# Canonical microRNA loss drives tumor development implicating therapeutic efficacy of enoxacin in angiosarcoma

**DOI:** 10.1101/2025.07.15.664801

**Authors:** Ant Murphy, Bozhi Liu, Annaleigh Benton, Lauren Gartenhaus, Samuel Hartzler, Madison E. Yates, Alejandra Romero Alzate, Nimod D. Janson, Shyaman Jayasundara, Sagar Utturkar, Nadia A. Lanman, Majid Kazemian, Jason A. Hanna

**Author notes:** Correspondence to: Jason A. Hanna, 201 S. University Street Department of Biological Science Purdue University, West Lafayette, IN 47906. These authors contributed equally (Ant Murphy, Bozhi Liu, Annaleigh Benton).

## Abstract

Angiosarcoma (AS) is a rare and aggressive tumor arising within the endothelium, characterized by a high metastatic rate and poor prognosis. Our prior work established that endothelial loss of *Dicer1*, a key enzyme in microRNA (miRNA) processing, drives AS formation in mice, indicating a tumor suppressive role for miRNAs in tumorigenesis. Here, we corroborated this hypothesis by generating a novel conditional knockout model targeting *Dgcr8*, a core component of the microprocessor complex required for pri-miRNA processing. Conditional deletion of *Dgcr8* phenocopies *Dicer1* loss, resulting in spontaneous AS formation and global loss of mature miRNAs. We further demonstrate that treatment with enoxacin (ENX), a repurposed antibiotic known to enhance miRNA processing, reduces viability, migration, and clonogenicity of AS cells. ENX increases the abundance of tumor-suppressive miRNAs and downregulates oncogenic pathways, including pathways related to cell cycle progression, angiogenesis, and cell migration. These results establish the essential role of miRNA biogenesis in suppressing AS and reveal a pharmacologically targetable vulnerability via ENX-mediated enhancement of miRNA expression in tumors.

## Introduction

Angiosarcoma (AS) is a rare tumor arising within the vascular or lymphatic endothelium and can occur in virtually any anatomic location. Comprising 2% of all soft tissue sarcomas in humans, AS is notoriously malignant with a metastatic rate of 44% and a five-year overall survival of 20-30%^1–3^. The molecular mechanisms underlying AS initiation and progression are not well understood; however, varied risk factors for AS have been identified, including: chronic lymphoedema, therapeutic radiation, UV exposure, exposure to carcinogenic compounds such as vinyl chloride, and familial genetic syndromes^4,5^. Treatments for AS are varied: surgery, radiotherapy, and chemotherapy are utilized depending on anatomic site, size, and presence of metastasis^1^. The diversity of AS necessitates the need for new models and tools to help understand the diverse mechanisms driving AS progression and how to address them pharmacologically.

We previously demonstrated that conditional deletion of *Dicer1* in endothelial cells and subsequent loss of mature microRNAs (miRNAs), leads to angiosarcoma development in a genetically engineered mouse model^6^. MiRNAs are regulatory RNAs which attenuate gene expression by either promoting degradation of specific mRNA transcripts, or by repressing their translation^7^. MiRNAs are important regulators in every cell type, and in virtually every cellular process^8^. They are transcribed by RNA Pol II as long transcripts, referred to as primary miRNAs (pri-miRNAs), which are then trimmed to smaller hairpins (pre-miRNAs) in the nucleus by the Microprocessor complex, consisting of DROSHA and DGCR8 (DiGeorge critical region 8) enzymes^9^. These RNA hairpins are exported to the cytoplasm via Exportin V where they are then processed to the final mature miRNA by a DICER1^10^. The mature miRNAs are then loaded into RNA-induced silencing complexes (RISC) containing Argonaut (Ago) family of proteins. The miRNA loaded RISC complex is then directed to target mRNAs based on partial miRNA-mRNA complementary^11^. This interaction leads to mRNA degradation and translational repression, ultimately downregulating the expression of the regulated genes.

Dysfunction of DGCR8 is well-known to be associated with Digeorge Syndrome; a rare genetic disorder in which a segment of chromosome 22 (including *DGCR8*) is deleted, causing a diverse set of symptoms including learning disabilities, congenital heart disease, weakened immune system, scoliosis, among others^12^. In addition to the wide range of disorders caused by loss of *DGCR8,* mutations and loss of heterozygosity of *DGCR8* have been associated with cancer^13,14^. Driver mutations in both microprocessor genes, *DGCR8* and *DROSHA,* have been identified in Wilms tumor and this leads to dysregulated miRNA processing^15–17^.

ENX, a fluoroquinolone antibiotic, has been discovered to enhance miRNA processing^18,19^. Although the precise mechanism is currently unclear, this effect is hypothesized to be accomplished through increasing the affinity between TRBP and pre-miRNAs, leading to enhanced processing by DICER1^20^. ENX promotes miRNA activity even in cancer cells with DICER1 downregulation, suggesting a potential therapeutic value by restoring miRNA activity in a biogenesis-impaired context^21^. Interestingly, studies that have evaluated miRNA abundance in ENX-treated cells indicate a similar number of decreased and increased miRNAs, yet ENX globally represses tumorigenesis and curiously tends to promote suppressive miRNA abundance^22^.

In our previous work, we demonstrated that conditional loss of *Dicer1* is sufficient to promote AS development using Cre recombinase under the control of the *adipocyte protein 2 (aP2)* promoter^23^. To determine if this phenotype is dependent on canonical miRNA loss and not due to other functions of DICER1, we have conditionally deleted *Dgcr8,* which processes miRNAs upstream of DICER1. We also evaluate the efficacy of ENX and its impact on gene and miRNA expression in AS cells.

## Results

### aP2-Cre;Dgcr8^Flox/Flox^ mice are viable

Following our recent observations that conditional *Dicer1* deletion in mice drives AS development^6,24^, we sought to determine if global miRNA loss was the driving force behind tumorigenesis or if non-canonical functions of DICER1 could be contributing. Therefore, we tested if the conditional deletion of *Dgcr8* phenocopied *Dicer1* loss and led to AS. We generated *aP2-Cre;Dgcr8^Flox/Flo**x**^* (AD8^cKO^) using the *aP2-Cre* mouse generated by Tang et al.^23^ and the Dgcr8^Flox/Flox^ mouse generated by Rao et al.^25^ (Fig. 1A). Recombination of *Dgcr8* by Cre is accomplished by lox P sites which flank exon 3, resulting in a nonfunctional transcript (Fig. 1B). *aP2-Cre* expression, and *Dgcr8* recombination is observed in adipose tissues as expected with high efficiency but with some retention of the Flox allele as previously observed (Fig. 1C)^6^.

**Figure 1.**
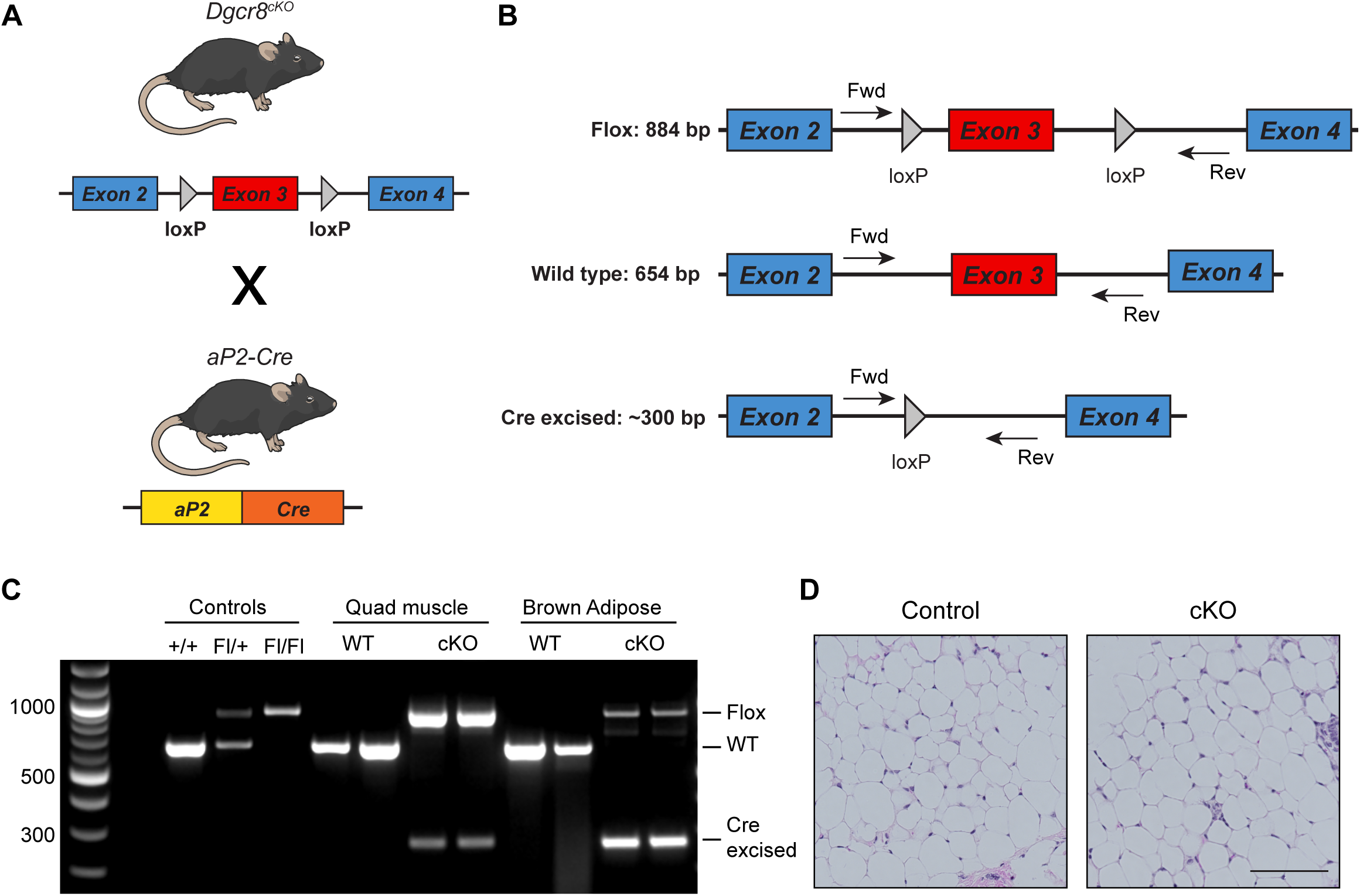
*Dgcr8* conditional deletion with *aP2-Cre*. **(A)** Graphic of *Dgcr8^cKO^* and *aP2-Cre* alleles and breeding scheme to generate *aP2-Cre;Dgcr8^cKO^*animals. Mouse illustration from NIAID NIH BIOART Source (bioart.niaid.nih.gov/bioart/279). **(B)** Map of primers for genotyping the Flox, Wild type, and Cre excised alleles by genomic PCR. **(C)** Genomic PCR of DNA from controls, quadricep muscle, or brown adipose tissue of wild type (WT) or *aP2-Cre;Dgcr8^Fl/Fl^*(cKO) animals. **(D)** Normal inguinal white adipose tissue H&E from a wild type control (Cre-, Dgcr8^Fl/Fl^) or Cre+, *aP2-Cre;Dgcr8^Fl/Fl^*(cKO) animal, scale bar 100µm.

Despite depletion of *Dgcr8* in adipocytes and endothelial cells, the AD8^cKO^ mice were viable and fertile with no apparent phenotypes or histologic findings to note in any adipose tissue including the inguinal white adipocyte fat pad from adult (8-week old) control and Dcgr8^cKO^ mice (Fig. 1D).

### Biallelic loss of *Dgcr8* drives angiosarcoma

Loss of a single *Dgcr8* allele (AD8^cHet^) is insufficient for tumorigenesis; however, *aP2-Cre;Dgcr8^Flox/Flox^* (*AD8^cKO^*) mice develop AS with a median tumor-free survival of 266 days (Fig. 2A). AS development was 100% penetrant and specific, with hemorrhagic AS tumors being the only tumors that were detected. A panel of endothelial-specific immunohistochemical staining for PECAM1, ERG, and Factor VIII demonstrates the endothelial nature of these tumors, consistent with AS (Fig. 2B). Tumors exhibit a high degree of recombination of *Dgcr8* with the non-recombined Flox allele, likely amplified from contaminating tumor stroma and normal blood cells (Fig. 2C, D). As expected, *AD8^cKO^*tumors have a significant reduction of *Dgcr8* transcript and mature miRNAs, reflecting the loss of a functional microprocessor complex (Fig. 2D, E).

**Figure 2.**
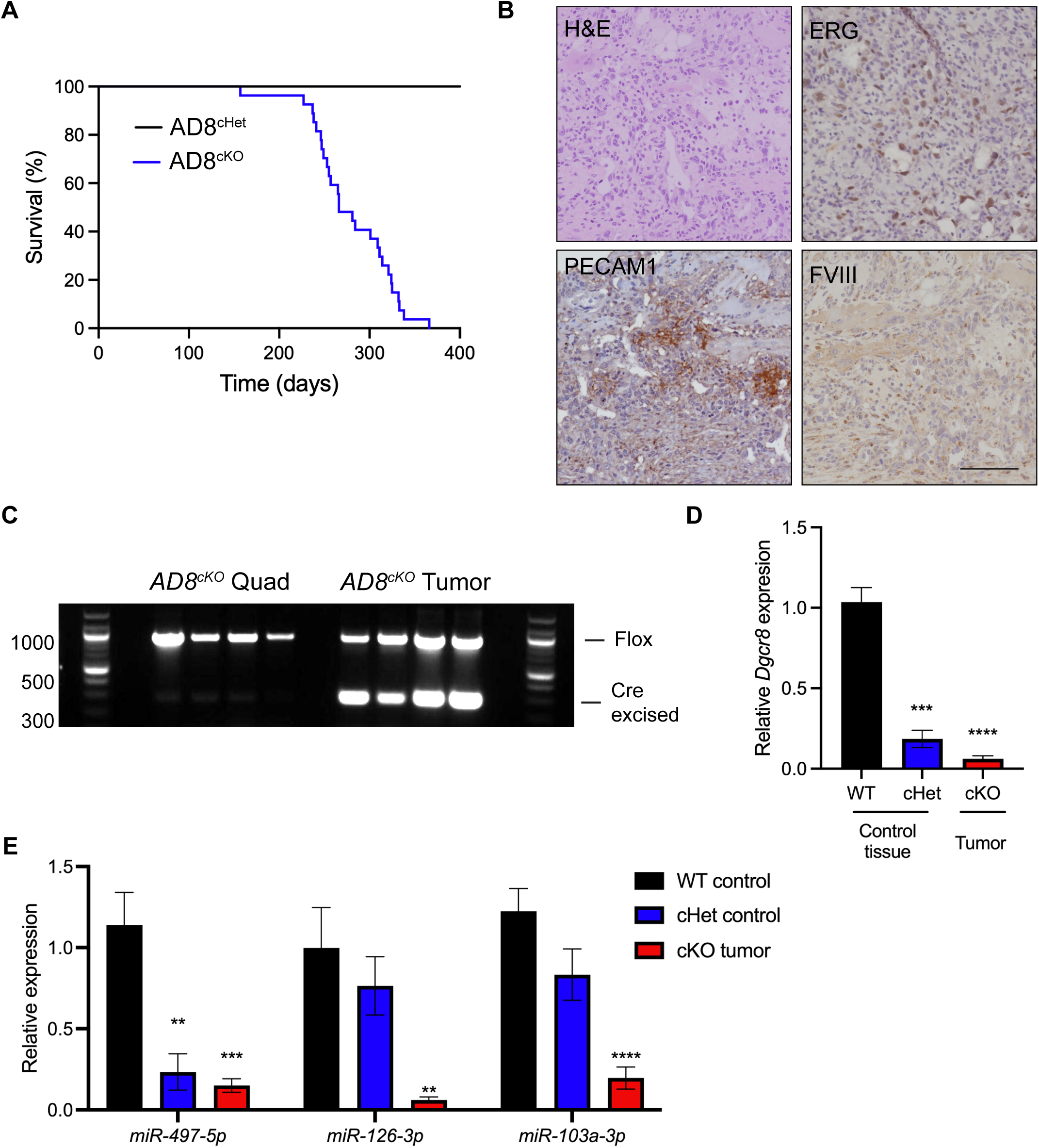
*Dgcr8* deletion leads to angiosarcoma development. **(A)** Kaplan-Meier tumor-free survival curve for *aP2-Cre;Dgcr8^cHet^*(AD8^cHet^, black, n=28) or *aP2-Cre;Dgcr8^cKO^* (AD8^cKO^, blue, n=27, median tumor free survival time of 266 days), log-rank p<0.0001**. (B)** H&E and IHC for ERG, PECAM1, or FACTOR VIII (FVIII) from an AD8^cKO^ tumor, scale bar 100µm. **(C)** Genomic PCR of DNA from AD8^cKO^ control quadricep (quad) muscle or AD8^cKO^ angiosarcomas (n=4). **(D)** Relative expression of *Dgcr8* by RT-PCR from WT and cHet control tissue (n=3) compared to cKO (AD8^cKO^) angiosarcomas (n=3). **(E)** Relative expression of indicated miRNAs by qRT-PCR from tissue as in (D), **p<0.01, ***p<0.001, ****p<0.0001.

Overall, biallelic loss of *Dgcr8* recapitulates the *Dicer1* deletion phenotype, suggesting that loss of canonical miRNA processing drives tumorigenesis and miRNAs are critical tumor suppressors in AS.

### Enoxacin treatment of AS cells reduces viability, clonogenic potential, and migratory ability

While most cells cannot survive miRNA loss, *Dgcr8* or *Dicer1* deletion in the aP2-Cre model promotes transformation and tumorigenesis, suggesting endothelial cells are uniquely susceptible to miRNA-loss-mediated transformation. Thus, we hypothesized that the restoration of miRNA expression may be particularly efficacious in AS. Treatment of murine AS cell lines with intact or reduced biogenesis machinery, SVR and ADC106, respectively, with 100 µM enoxacin (ENX) resulted in a significant reduction in cell viability and a significant increase in Caspase 3/7 activity, indicating an increase in apoptosis (Fig. 3A, B). This effect was more pronounced in AS cells since ENX only subtly reduced normal HTHM (HUVEC cells immortalized with hTERT and MYC) but did not induce apoptosis (Fig. 3A, B). Furthermore, ENX-treated AS cells displayed a significantly lower population doubling time over a period of 10 days (Fig 3C). Using a standard colony formation assay, we observed a striking reduction in clonogenic ability (Fig 3D, E). Finally, ENX treatment resulted in a reduction in migratory ability of AS cells in standard transwell migration assays (Fig 3F, G).

**Figure 3.**
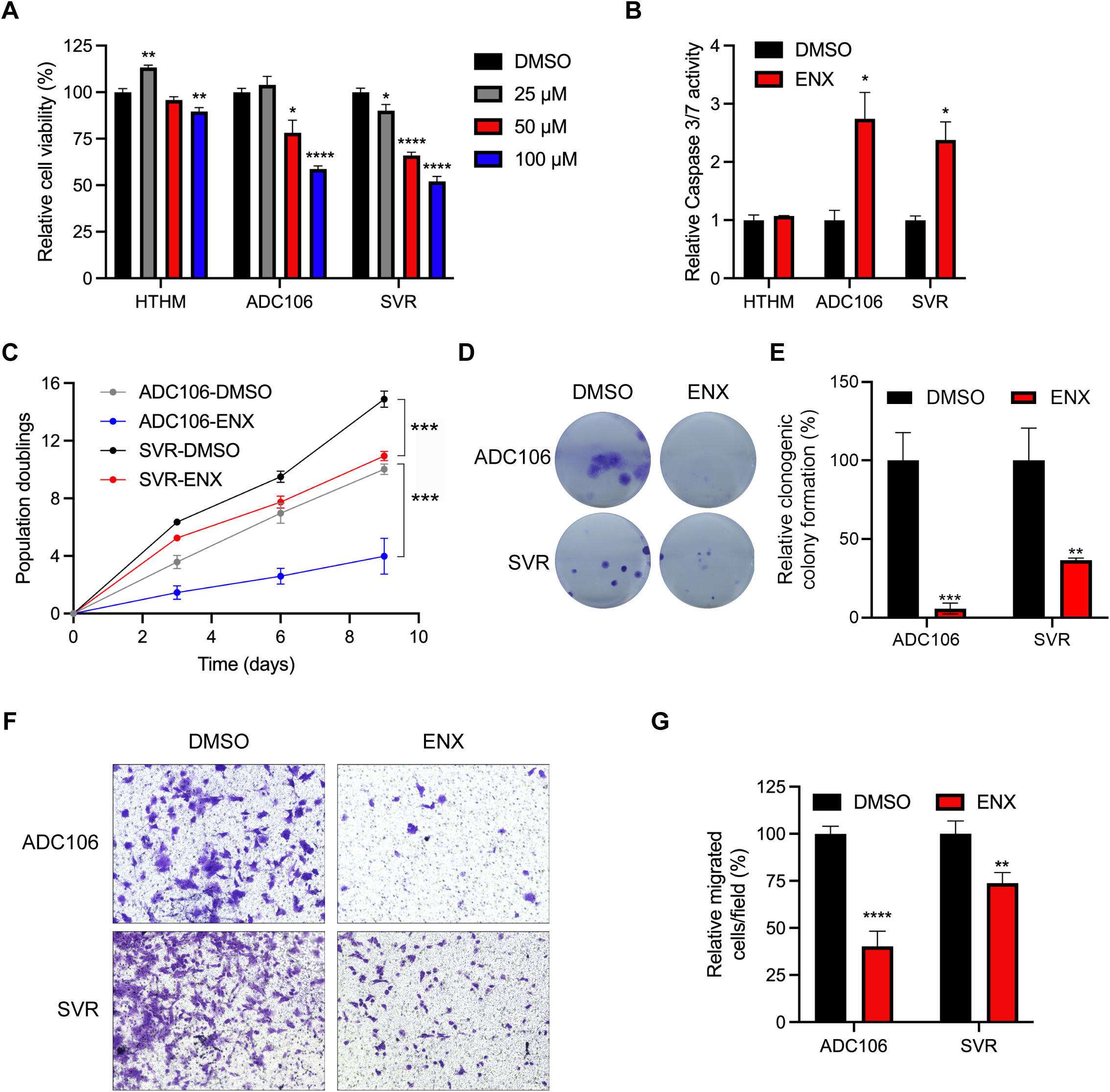
Enoxacin reduces angiosarcoma proliferation. **(A)** Relative cell viability via Cell Titer Glo in HTHM, ADC106, and SVR cells treated with indicated dose of enoxacin (ENX) for 72 hours. **(B)** Relative Caspase 3/7 activity in ADC106, SVR, and HTHM cells treated with DMSO or 100 µM ENX for 48 hours. **(C)** Population doubling assays in ADC106 and SVR cells treated with DMSO or 100µm ENX. **(D)** Representative images and **(E)** quantification of clonogenic colony formation in cells treated with DMSO or 100 µM ENX. **(F)** Representative image and **(G)** quantification of migrated cells in transwell migration assays in cells treated with DMSO or 100 µM ENX, data presented as the mean of four biological replicate experiments with standard error of mean (SEM), *p<0.05, **p<0.01, ***p<0.001, ****p<0.0001.

### Enoxacin treatment leads to increased miRNA expression and transcriptional changes

To determine whether ENX enhances miRNA biogenesis as hypothesized, we first tested its effects using a miR-497 luciferase reporter assay. This reporter, which we previously generated, contains a perfect miR-497 site inserted into the 3‘UTR of *Renilla* luciferase^26^. The construct was transiently transfected into HEK-293T cells, which were subsequently treated with ENX or DMSO control. As expected, a significant reduction in luciferase activity demonstrated that ENX likely promotes the processing and/or abundance of miR-497 (Supplemental Fig.S1A).

To determine global effects on miRNA abundance and mRNA expression, we conducted small RNA-seq and mRNA-seq on SVR and ADC106 cells treated with DMSO or ENX for 72 hours. To evaluate whether ENX alters miRNA processing rather than transcription, we first examined the expression of several intronic miRNAs and their host genes (Fig. 4A). Increased miRNA abundance was observed without corresponding increases in host gene transcript levels, implicating increased miRNA biogenesis as the driver of miRNA upregulation, not transcription (Fig. 4B). We also evaluated other post-transcriptional mechanisms such as LIN28B regulation of the let-7 family of miRNAs. LIN28B represses the biogenesis of let-7 by inhibiting its processing by DROSHA/DGCR8 in the nucleus^27–29^. Therefore, decreased *Lin28b* expression could increase let-7 abundance^30^. We observed no decrease in *Lin28b expression*, while many let-7 family miRNAs were significantly increased in abundance (Fig.4D). These finding suggest that let-7 upregulation is not mediated by *Lin28b* downregulation, but rather by other post-transcriptional mechanisms, consistent with the hypothesis that ENX enhances miRNA processing.

**Figure 4.**
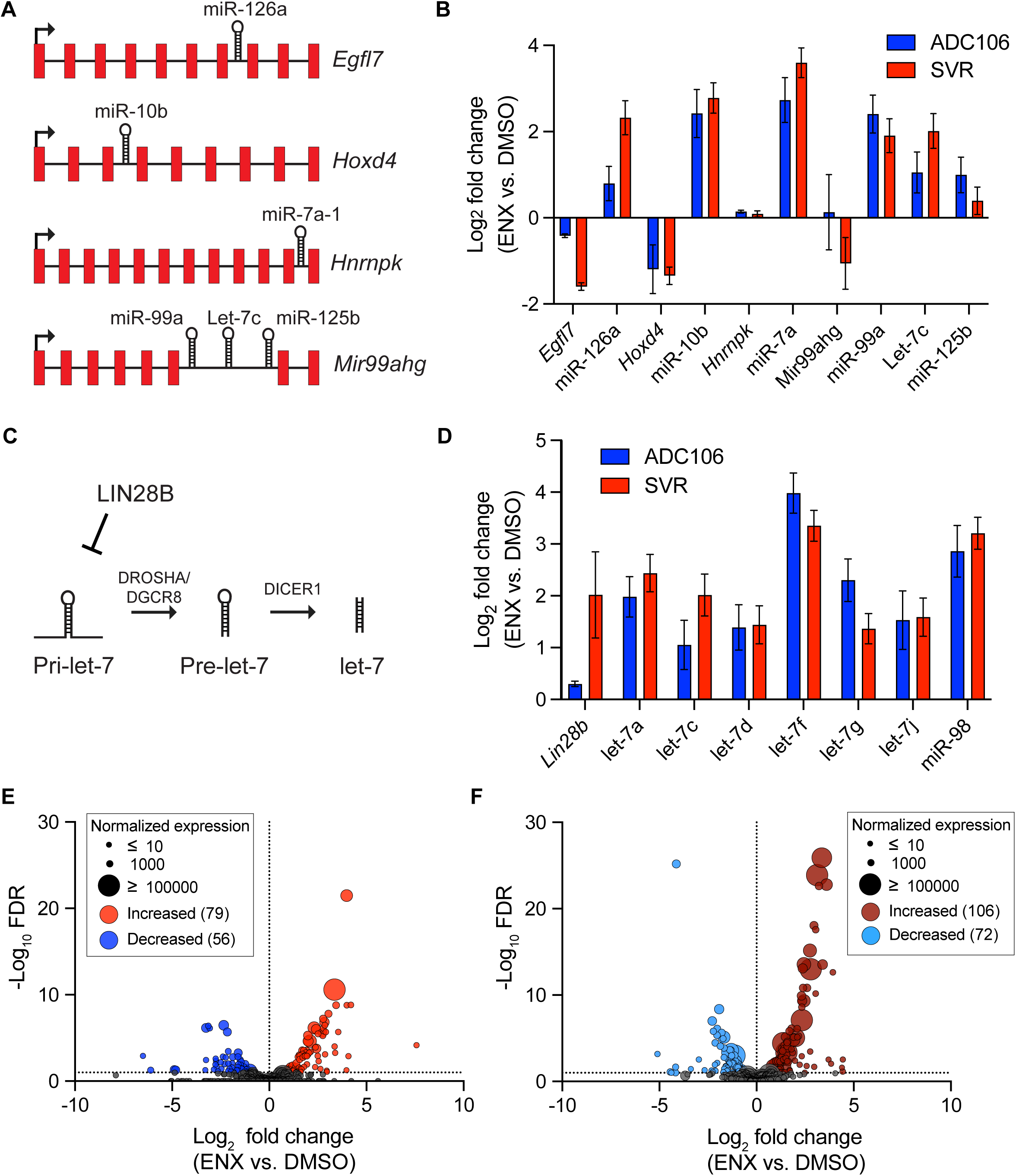
Enoxacin enhances miRNA biogenesis. **(A)** Graphic of intronic miRNAs and their host genes. **(B)** Log_2_ fold change in expression comparing enoxacin (ENX) vs DMSO treated ADC106 (blue) or SVR (red) of intronic miRNAs and host gene transcripts by RNA-seq. **(C)** Graphic of LIN28B regulation of let-7 miRNAs. **(D)** Log_2_ fold change in expression of *Lin28b* and let-7 family miRNAs by RNA-seq. **(E)** Volcano plot of the-log_10_ FDR vs the log_2_ fold change in miRNA abundance of ADC106 cells or **(F)** SVR cells treated with DMSO or 100 µM ENX for 72 hours. Differentially expressed miRNAs with increased or decreased abundance with FDR < 0.05 and log_2_ fold change > 1 and <-1 as indicated.

We observed 135 differentially expressed miRNAs (absolute log_2_ fold change > 0.5, p value < 0.05) in ADC106 cells treated with ENX compared to DMSO-treated control cells, with a slight majority being upregulated (79 increased vs. 56 decreased) (Fig. 4E, Supplemental Fig. S1B). Similarly, ENX-treated SVR cells had 178 miRNAs differentially expressed (106 increased vs. 72 decreased) (Fig. 4F, Supplemental Fig. S1C). Notably, SVR and ADC106 cells exhibited distinct but overlapping miRNA expression profiles (Supplemental Fig. S1D-F and S2), with many differentially expressed miRNAs shared between the two cell lines. Notably, the 50 commonly upregulated miRNAs include several known tumor suppressors, such as let-7 family, miR-206, miR-181, miR-146a, miR-335, and miR-126. Additionally, members of the miR-15/16/497/195/424 family, which we have previously shown to be potent tumor-suppressing miRNAs^26^, were also increased in the ADC106 cells.

To determine whether ENX also influences mRNA expression more broadly, we performed RNA-seq on SVR and ADC106 cells treated with ENX or DMSO for 72 hours. We identified 1,689 and 3,733 differentially expressed mRNAs (absolute log_2_ fold change > 0.5, FDR < 0.05) in ADC106 and SVR cells, respectively, compared to DMSO-treated controls, with a slight majority being downregulated (Supplemental Fig. S3A-C). To gain insight into the biological processes affected by ENX treatment, we next performed gene ontology (GO) analysis using DAVID. GO analysis of the upregulated genes (FDR < 0.05, log_2_ fold change > 0.5) identified terms such as programmed cell death, negative regulation of cell proliferation, cell differentiation, and ribosome biogenesis (Fig. 5A, B, Supplemental Tables S1, S2). Notably, several pro-apoptotic, growth arrest, and differentiation genes such as *Gadd45a*, *Rb1*, *Rassf6*, Wwox, *Dapk1*, *Rasa3*, *Btg3*, *Steap1*, *Klf4*, and *Deptor* were upregulated in both cell lines (Supplemental Fig. S3D), whereas others, such as *Cdkn1a*, *Rb1*, *Fas*, *Rassf5*, *Chd5*, and *Plk3,* were increased in only one of the two. In contrast, among the GO pathways associated with the downregulated genes, we observed common terms such as cell cycle, angiogenesis, and vascular development (Fig. 5A-B, Supplemental Tables S3, S4). Angiogenesis-related genes such as *Angpt1*, *Egfl7*, *Flt4*, *Notch4*, and *Tek* were consistently downregulated in both cell lines (Supplemental Fig. S3E). We also conducted Gene Set Enrichment analysis (GSEA) and evaluated the top 5 enriched Hallmark pathways in both cell lines based on absolute normalized enrichment scores (NES). The cell cycle-related pathway “E2F Targets” was significantly suppressed in both cell lines (Fig. 5C, D, Supplemental Fig. S4A-C). In SVR cells, pathways involved in cell stress response and inflammation were also suppressed while IL6-JAK-STAT3 signaling was activated. Interestingly TNFIZI signaling, oxidative phosphorylation, and the P53 pathways were enriched in ADC106 cells, whereas E2F targets and G2/M checkpoint pathways were decreased. Collectively, these data indicate that ENX broadly impacts mRNA expression.

**Figure 5.**
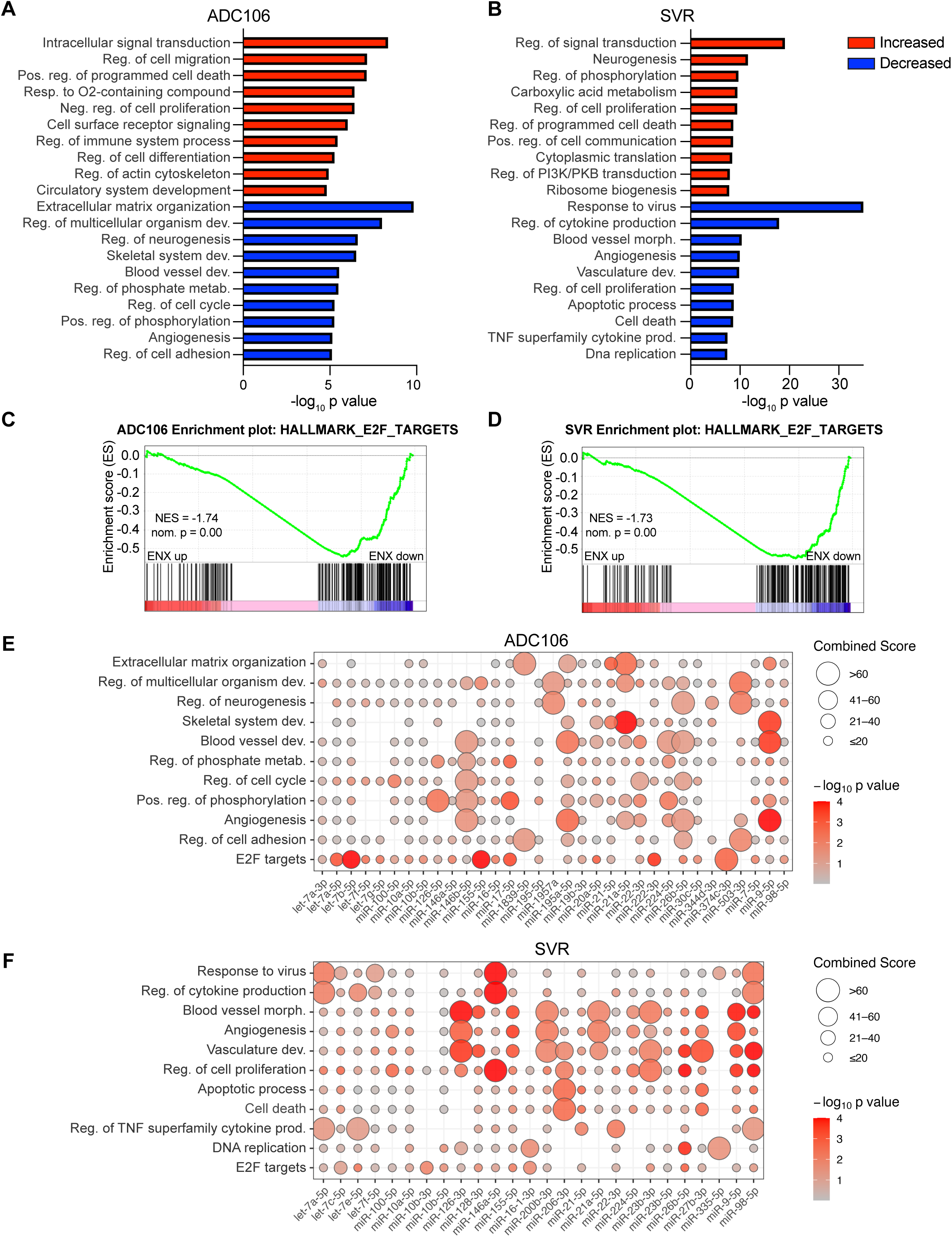
Enoxacin alters mRNA expression. (A-B) DAVID Gene ontology analyses of significantly upregulated (red) and downregulated (blue) genes in ADC106 (A) or SVR (B) cells. **(C-D)** GSEA of the hallmark pathway “E2F targets” in the ADC106 (C) and SVR (D) cells. Normalized enrichment scores (NES) and nominal p values are shown. **(E-F)** Enrichment plot of potential miRNA-regulated genes with the downregulated genes (FDR < 0.05, log_2_ fold change <-0.5) within the top oncogenic pathways based on DAVID Gene ontology or the overlapping Hallmark GSEA terms in ADC106 (E) or SVR cells (F) treated with 100 µm ENX or DMSO.

### Enhanced miRNA expression regulates oncogenic pathways

We hypothesize that the anticancer properties of ENX can be attributed to the increased abundance of mature tumor-suppressing miRNAs. To test this, we evaluated the enrichment of miRNA-regulated genes within the downregulated DAVID GO and E2F Targets GSEA pathways using miRTarBase for experimentally validated miRNA-target genes^31^. Indeed, each suppressed pathway contained at least one statistically significant miRNA with enriched target genes (Fig. 5E, F, Supplemental Tables S5 and S6). Some pathways, such as the E2F targets, cell proliferation, and angiogenesis pathways, contained multiple miRNAs with enriched target genes. Interestingly, different miRNA target genes were enriched in each cell line. For example, in ADC106 cells, downregulated let-7 target genes were involved in cell cycle progression and E2F targets. In contrast, in SVR cells let-7 target genes involved in cytokine production were more prominently downregulated. Consistently, miR-9 target genes in both cell lines were associated with angiogenesis, while miR-146 target genes were linked to both cell proliferation and angiogenesis.

To identify critical pathways in AS potentially regulated by miRNAs we determined the intersection between genes downregulated by ENX in both cell lines and the genes upregulated in *Dicer1^cKO^* AS, where miRNA expression is lost (Fig. 6A)^6^. Gene ontology of this shared gene set revealed enrichment for pathways involved in cell cycle regulation, angiogenesis, and migration (Fig. 6B). Finally, miRTarBase analysis of the 465 genes (limited to just the 50 miRNAs increased in abundance in both cell lines) revealed significant enrichment of well-known tumor suppressing miRNAs as potential regulators of many of these oncogenic genes including let-7 family (miR-98, let-7c, let-7a, let-7e, let-7d, and let-7g), miR-155-5p, and miR-126a among others (Fig. 6C). These data suggest that ENX treatment exerts its anti-tumor effects by restoring miRNA-mediated suppression of oncogenic transcripts and thereby inhibits key pathways such as cell proliferation, angiogenesis, and migration (Fig. 6D).

**Figure 6.**
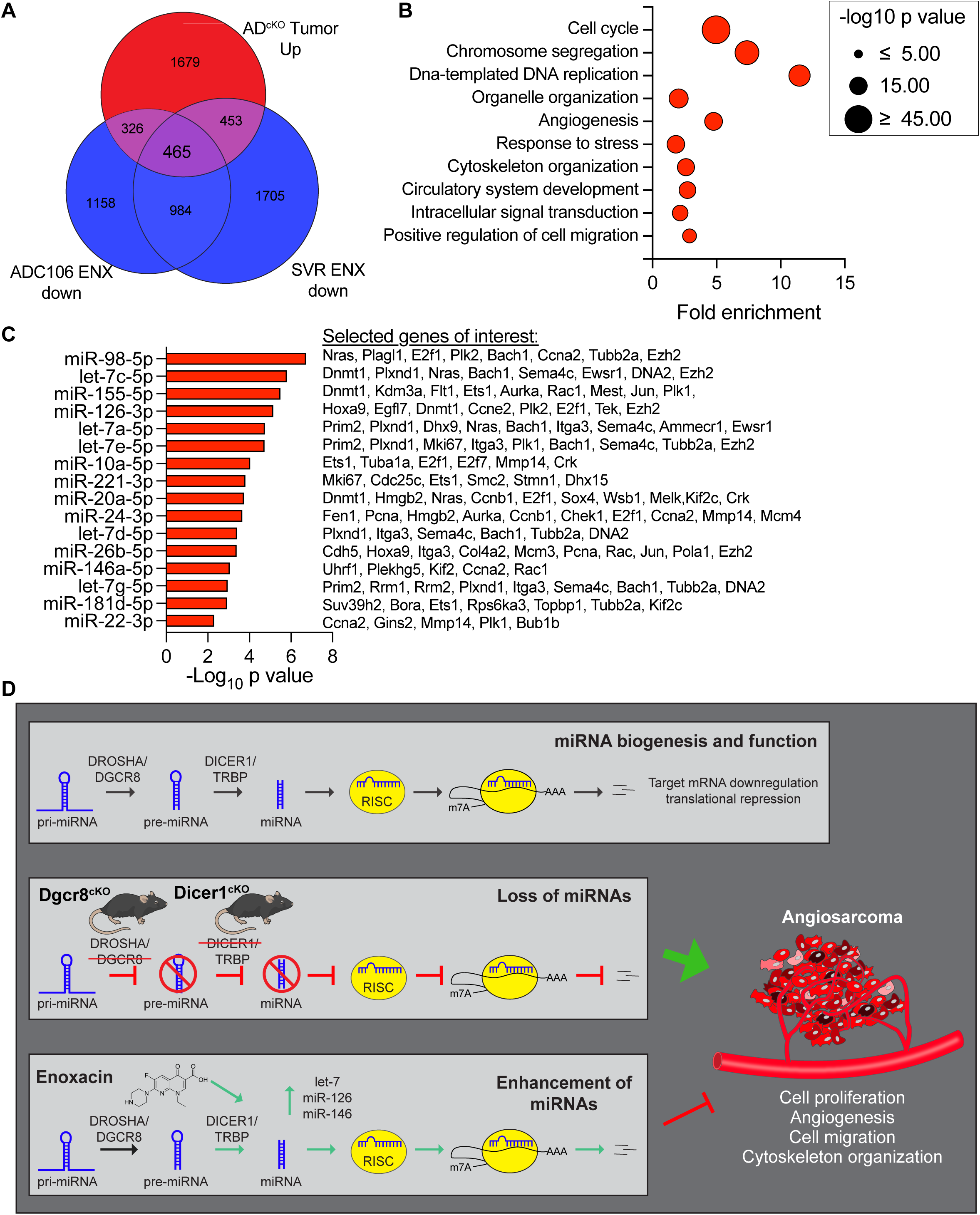
Commonly enriched miRNAs regulate cell cycle, angiogenesis, and cell migration pathways in angiosarcoma. **(A)** Euler diagram of the overlapping genes that are increased in expression in *aP2-Cre;Dicer1^Fl/Fl^* (AD^cKO^, from GSE85834). tumors compared to normal aorta (FDR < 0.05, log_2_ fold change > 0.5) and all genes with decreased expression in ADC106 or SVR cells (FDR < 0.0). **(B)** DAVID gene ontology analysis of enriched terms within the 465 overlapping genes from (A). **(C)** The top miRNAs with increased abundance in common between ADC106 and SVR cells treated with ENX with enrichment of target genes among the overlapping genes from (A). All miRNAs with significant (FDR < 0.05) enrichment of target genes plotted. Selected oncogenic genes experimentally validated to be regulated by miRNAs indicated. **(D)** Graphical abstract illustrating normal miRNA biogenesis and function of miRNAs. Conditional *Dgcr8* or *Dicer1* deletion in aP2-Cre expressing cells disrupts canonical miRNA biogenesis, leading to angiosarcoma development in mice. This highlights the important tumor-suppressing role of miRNAs in angiosarcoma. Pharmacologic repurposing the antibiotic enoxacin enhances the expression of several tumor suppressing miRNAs and demonstrates anticancer properties in angiosarcoma by repressing oncogenic pathways such as cell proliferation, angiogenesis, cell migration, and cytoskeleton organization. Mouse illustration from NIAID NIH BIOART Source (bioart.niaid.nih.gov/bioart/279).

## Discussion

Most human tumors exhibit a global repression of miRNAs, indicating the importance of downregulating tumor-suppressing miRNAs in cancer^32–35^. In this study, we demonstrated that *Dgcr8* loss in *aP2-Cre* expressing cells phenocopies the AS development driven by *Dicer1* deletion, indicating global miRNA repression as the molecular driver of tumorigenesis. *Dicer1* downregulation promotes tumor development in many mouse models of cancer^36–38^. We have previously shown that *Dicer1* is a classic tumor suppressor in endothelial cells, in which loss of both alleles leads to tumor development^6,24^. In contrast, in most other cell types, DICER1 functions as a haploinsufficient tumor suppressor, in which loss of both alleles is detrimental to tumor development and cell survival^36,37^. We find that endothelial cells not only survive *Dicer1* or *Dgcr8* biallelic deletion, but this is transformative and leads to aggressive AS development. This indicates that endothelial cells are uniquely sensitive to miRNA loss-mediated transformation.

Although individual miRNAs may have tumor suppressive or oncogenic functions, global suppression of miRNAs in endothelial cells drives tumorigenesis. This underscores the critical tumor-suppressive role of miRNA regulation of gene expression in AS. Based on our observations of AS development with miRNA loss, we hypothesized that AS cells may be sensitive to miRNA restoration-based therapies. Thus, we tested Enoxacin (ENX), a fluoroquinolone antibiotic, previously shown to enhance miRNA biogenesis^18,20^. ENX is thought to act by enhancing interactions between TRBP and pre-miRNA hairpins, thereby increasing DICER1-mediated processing of the hairpins^19,20^. However, this precise mechanism is still unclear. Genetic studies indicate DICER1 is required for ENX effects, but some studies call into question the role of TRBP^39^. Nonetheless, consistent with the tumor suppressive effects of miRNAs in AS, we found that AS cell lines treated with ENX were less proliferative and migratory and exhibited increased apoptosis.

By small RNA-seq, we determined that ENX induced significant changes in the expression of many miRNAs, while a slight majority of miRNAs were enhanced with ENX in both cell lines, a significant number were also decreased as has been previously reported^19,20,40–42^.

Like others have speculated, we hypothesize these downregulated miRNAs could be due to secondary consequences or pleiotropic effects of ENX altering the expression or stability of these miRNAs. It is also possible ENX alters the affinity of DICER1/TRBP for some miRNAs, however this has not been experimentally tested.

We identified several well-characterized tumor suppressive miRNAs with significantly increased abundance in both ADC106 and SVR cells. Notably, multiple members of the let-7 family were increased. Let-7 has widely been implicated as a tumor suppressor in many cancer types and is known to regulate several oncogenes and mediate cell proliferation, differentiation, and apoptosis^33^. miR-126 was also significantly increased with ENX; this endothelial enriched miRNA has been shown to maintain vascular integrity and inhibit tumor angiogenesis, further supporting its potential as a tumor suppressor in AS. Importantly, we also observe significant increased expression of miR-16-5p and miR-195-5p, members of the miR-15/16/497/424 family, in the ADC106 cells and a subtle increase in miR-16 in the SVR cells. We and others previously identified this family as a potent regulator of cell proliferation, apoptosis, and cell migration in AS^26,43^. Future studies on the individual and combined functions and mechanisms of these miRNAs will be important.

In addition to its anticancer effects^20,44^, ENX has also been investigated as an antiviral^40^, as a treatment for ALS^45,46^, to prevent obesity and extend lifespan^42,47^, and in the context of inhibiting osteoclasts in bone disease^48,49^. As an antibiotic, ENX was previously approved by the FDA, but is no longer widely used due to better efficacy with other drugs and potential side effects. However, due to its established clinical history, ENX may have a relatively straightforward path as an anticancer, either as a monotherapy or in combination with other therapies.

## Materials and Methods

### Mouse studies

All mouse strains were previously generated and described including: *aP2-Cre* (*Tg(Fabp4-Cre)1Jmgr*), MGI: 5300925)^23^, *Dgcr8^Flox^* (B6N.Cg-Dgcr8tm1.1Blel/Mmjax), MGI: 4458072). All mice were maintained on a mixed genetic background. Therefore, littermate controls were used for each study. Mice were fed and watered *ad libitum* in a facility with maintained humidity and ambient temperature with 12-hour light/dark cycles. Genomic DNA was isolated from tissue using the DNeasy Blood and Tissue Kit (69504, Qiagen). *aP2-Cre* and *Dgcr8* genotypes were determined as described previously and primers detailed in Supplemental Table S7^23,25^.

All experiments involving animal studies were reviewed and approved by the Purdue University Institutional Animal Care and Use Committee (protocol #1908001941).

### Cell Culture

Cells were obtained from the following sources: SVR (ATCC, Cat# CRL-2280, RRID:CVCL_6455), ADC106 (derived from our lab as previously described^24^), HTHM (HUVEC cells immortalized with large T (genomic) antigen, hTERT, and MYC)^50^, and HEK-293T (RRID:CVCL_0063, M. Kazemian, Purdue University). The SVR, ADC106, and HEK-293T cells were maintained in DMEM (SH30243, Hyclone) with 10% FBS (SH30910.03, Hyclone), 1× antibiotic-antimycotic penicillin, streptomycin, and amphotericin B (PSA) (A5955, Sigma-Aldrich, St. Louis, MO, USA). The HTHM cells were maintained in VascuLife VEGF endothelial media (Lifeline cell technology, LM-0002, Frederick, MD, USA), corresponding supplementary kit (Lifeline cell technology, LS-1020), and 1x PSA. All cells incubated at 37 °C in 5% CO_2_. Cell lines were passaged no more than 15 times past the initial thaw, and mycoplasma testing is performed 3 times annually, with each experiment being performed within 4 months of the last mycoplasma testing.

Cells were treated with DMSO or enoxacin (ENX) (Cat #J61912.06; Thermo Scientific, Waltham, MA, USA). Cell viability was determined using the Cell Titer Glo Assay (G7570, Promega, Madison, WI, USA) and apoptosis was determined using the Caspase 3/7 Glo Assay (G8090, Promega) measuring luminescence with a BioTek Synergy 2 (BioTek/Agilent, Winooski, VT, USA, RRID:SCR_020536). Population doubling assay was performed as previously described^51^. In brief, 50,000 cells were seeded, and every 3 days, cells were lifted and counted using the Luna II Automated Cell Counter (Logos Biosystems, Annandale, VA, USA). Then 50,000 cells were reseeded for continued culture. Colony formation was performed by seeding 50 or 100 cells per well in 6-well plates and culturing them under standard conditions. When visible colonies had formed, cells were stained with crystal violet for 15 minutes. Excess stain was rinsed off with distilled water, and plates were air-dried. Transwell assays were performed with transwell permeable supports (3422, Corning, Glendale, AZ, USA). Cells were suspended in serum free DMEM, and DMEM supplemented with 10% FBS was used as a chemoattractant. Non-migrated cells were removed from the inner chamber with a cotton swab, migrated cells were crystal violet stained, imaged, and quantified. At least three independent fields per sample were used for quantification of images. Dual luciferase assays were performed by seeding HEK-293T cells in 48-well plates as described^52^. The following day, cells were treated with DMSO or 100 µM ENX. On the next day, cells were transfected with FuGENE6 (Cat# E2691; Promega) 5Dng of psiCHECK2 reporter plasmids (empty or a miR-497 sensor in which the Renilla luciferase 3’UTR was modified to include a perfectly complementary miR-497 site as described previously^26^). 72 hours post transfection, cells were lysed in passive lysis buffer, and luciferase activity was measured using the Dual-Luciferase Reporter Assay System (Cat# E1910; Promega) on a SpectraMax multi-mode plate reader (Molecular Devices, RRID:SCR_023920). Renilla luciferase activity was normalized to Firefly luciferase activity to control for transfection efficiency and analyzed as relative luciferase activity compared to the DMSO treated empty control.

### Histology, and immunohistochemistry

Hematoxylin and eosin (H&E) histology and immunohistochemistry (IHC) were performed following standard protocols and antibodies against ERG (MA5-32036, Thermo Fisher Scientific), Factor VIII (ab203590, Abcam, Waltham, MA, USA), and PECAM1 (MA5-37858, Thermo Fisher Scientific). The antibodies and staining protocols details are provided in Supplemental Table S8.

### RNA and gene expression

Total RNA was prepared using the *Quick*-RNA Miniprep Kit (R1054, Zymo, Irvine, CA, USA) according to the manufacturer’s instructions. cDNA was generated using High-Capacity RNA to cDNA kit (4387406, Applied Biosystems, Waltham, MA, USA). Relative expression by qRT-PCR was quantified using the delta-delta CT method normalized to *Gapdh* using primers described in Supplemental Table S7. For miRNA expression, the miRCURY cDNA kit (339340, Qiagen, Germantown, MD, USA) was followed with qRT-PCR using LNA miRCURY primers detailed in Supplemental Table S7. MiRNA expression was normalized to snRNA U6. RNA-seq was conducted on total RNA from ADC106 or SVR cells treated with DMSO or 100 µM ENX for 72 hours. mRNA (poly-A enrichment) and small RNA library prep and sequencing were performed by Novogene on an Illumina NovaSeq platform. Differential gene and miRNA expression were analyzed as previously described^53^. In brief, for mRNA-seq, raw paired-end sequencing data were processed and quantified using RSEM (v1.3.1) with default parameters, and Bowtie2 as the aligner^54,55^. The Bowtie2 reference index was generated using ‘rsem-prepare-reference’ with the GRCm38 transcriptome and RefSeq gene annotations obtained from the UCSC Genome Browser. Differential gene expression analysis was performed using DESeq2 (v1.42.1)^56^. For the small RNA-seq, raw reads were cleaned (adapter trimming, quality trimming Phred quality score >30) using the Fastp toolkit (version 0.23.2)^57^. miRNA quantification was performed with QuickmiRSeq tool version 1.0^58^. Filtered counts from QuickmiRSeq were used for differential expression analysis with DEseq2 (v1.42.1). Previously published transcriptomic data from *aP2-Cre;Dicer1^Fl/Fl^* (AD^cKO^) from GSE85834^6^. Gene ontology analysis was conducted on the differentially expressed genes using the Database for Annotation, Visualization, and Integrated Discovery (DAVID)^59^. Classification of terms were analyzed based on biological process terms (BP_FAT) with the exclusion of redundant and general terms. Gene Set Enrichment Analysis (GSEA^60,61^) performed against the Hallmarks Gene Set. Enrichment of experimentally validated miRNA-targets gene regulation identified by Enrichr^62^ and the miRTarBase^63^ database.

## Statistics

Statistical analyses were performed using Prism Version 9 (Graph Pad Software, Inc., San Diego, CA, USA). All results are expressed as the mean ± SD unless stated otherwise. Pairwise comparisons were performed with a two-tailed, unpaired Student’s t-test. Significance cutoff at p values < 0.05 was considered significant.

## Acknowledgements

We thank Megan Cohen and the Purdue Histology Research Laboratory, a core facility of the NIH-funded Indiana Clinical and Translational Science Institute. We are supported by grants from the Purdue Institute for Cancer Research (P30CA023168), the Walther Cancer Foundation, the Purdue Institute for Cancer Research Summer Undergraduate Research Program, the Purdue Institute for Cancer Research Jim and Diann Robbers Cancer Research Grant for New Investigators Award (JAH), the extramural research programs of the NIH (R35GM138283 to MK) and the CDMRP Department of Defense Rare Cancers Research Program (HT94252410861 to JAH).

## Data and resource availability

All relevant data and resources can be found within the article and its supplementary data. The RNA-sequencing data are deposited in the Gene Expression Omnibus (GEO) under accession numbers GSE298851 and GSE298666.

## Author contributions statement

Conceptualization: JAH; Methodology: AM, AB, BL, SJ, SU; Formal Analysis: SJ, SU, NL, MK, JAH; Investigation: AM, AB, BL, LG, SH, MY, ARA, NDJ, SJ, SU, JAH; Writing original draft: AM, BL, JAH; Writing, reviewing, editing: AM, AB, BL, LG,SH, MY, ARA, NDJ, SJ, SU, NL, MK, JAH; Visualization: SJ, SU, NL, MK, JAH; Supervision: NL, MK, JAH; Funding acquisition: JAH.

