## Supplemental Figures for "Canonical microRNA loss drives tumor development implicating therapeutic efficacy of enoxacin in angiosarcoma"

### Supplemental Information Content

#### Supplemental Figures

**Supplemental Figure S1**. miRNA alterations with ENX.
**Supplemental Figure S2**. Heatmap of altered miRNA expression with enoxacin.
**Supplemental Figure S3.** mRNA expression changes with enoxacin.
**Supplemental Figure S4**. Gene Set Enrichment Analysis (GSEA) in cells treated with enoxacin.

#### Supplemental Tables

**Supplemental Table S1.** Gene ontology analysis of enoxacin upregulated genes in ADC106 cells.
**Supplemental Table S2.** Gene ontology analysis of enoxacin upregulated genes in SVR cells.
**Supplemental Table S3.** Gene ontology analysis of enoxacin downregulated genes in ADC106 cells.
**Supplemental Table S4**. Gene ontology analysis of enoxacin downregulated genes in SVR cells.
**Supplemental Table S5.** Predicted miRNA regulated genes in top downregulated pathways for miRNAs with increased abundance in ADC106 cells treated with 100μM ENX.
**Supplemental Table S6.** Predicted miRNA regulated genes in top downregulated pathways for miRNAs with increased abundance in SVR cells treated with 100μM ENX.
**Supplemental Table S7.** Primers used for qRT-PCR and genotyping.
**Supplemental Table S8.** Antibodies used for IHC.

**Supplemental figure legends**

**Supplemental Figure S1. miRNA alterations with ENX. (A)** Relative luciferase activity of miR-497 sensor reporter in HEK293T cells transfected with psiCHECK2-miR-497 sensor reporter and treated with DMSO or 100 µM ENX for 48 hours. **(B)** MA plot of small RNA-seq data in ADC106 or **(C)** SVR cells ($log_2$ fold change of ENX vs. DMSO). A base mean cut-off of 10 applied as the minimum expression level. **(D)** Combined volcano plot of differentially expressed miRNAs in the ADC106 and SVR cells. **(E)** Euler diagram of the common miRNAs increased in abundance with FDR < 0.05 and $log_2$ fold change > 0.5. Euler diagram with number of miRNAs with increased abundance in ENX vs. DMSO with FDR < 0.05 in ADC106 and SVR cells. **(F)** Multivariable bubble plot of the 50 common miRNAs of increased abundance based on the mean normalized expression (base mean) and differential expression ($log_2$ fold change) in ADC106 (blue) and SVR (red) cells.

**Supplemental Figure S2. Heatmap of altered miRNA expression with enoxacin.** Heatmap of the commonly dysregulated miRNAs by small RNA-seq in the ADC106 and SVR cells treated with ENX or DMSO for 72 hours.

**Supplemental Figure S3. mRNA expression changes with enoxacin. (A-B)** Volcano plots showing $-log_{10}$ FDR versus $log_2$ fold change in mRNA expression in ADC106 (A) or SVR (B) cells treated with enoxacin (ENX) or DMSO for 72 hours. Genes significantly downregulated (FDR < 0.05, $log_2$ fold change < -0.5; blue) or upregulated (FDR < 0.05, $log_2$ fold change > 0.5; red) are highlighted. **(C)** Gene expression data from RNA-seq were z-scored normalized. Hierarchical clustering of genes (rows) and samples (columns) of consistent direction and magnitude of fold change in both cell lines (mean $|log_2FC| > 0.5$) included. Color scale represents relative expression (z-score). **(D)** $Log_2$ fold change in expression of upregulated genes in ADC106 (blue) or SVR (red) cells treated with DMSO or ENX. **(E)** Expression of downregulated angiogenic and endothelial genes in cells as in (D).

**Supplemental Figure S4. Gene Set Enrichment Analysis (GSEA) in cells treated with enoxacin. (A)** Bar graphs of the Normalized Enrichment (NES) scores of the top pathways from GSEA analysis of the differentially expressed genes in ADC106 or **(B)** SVR cells**. (C)** Differential expression ($log_2$ fold change) of selected genes from the cell cycle and E2F targets gene sets in ADC106 (blue) or SVR (red) cells.

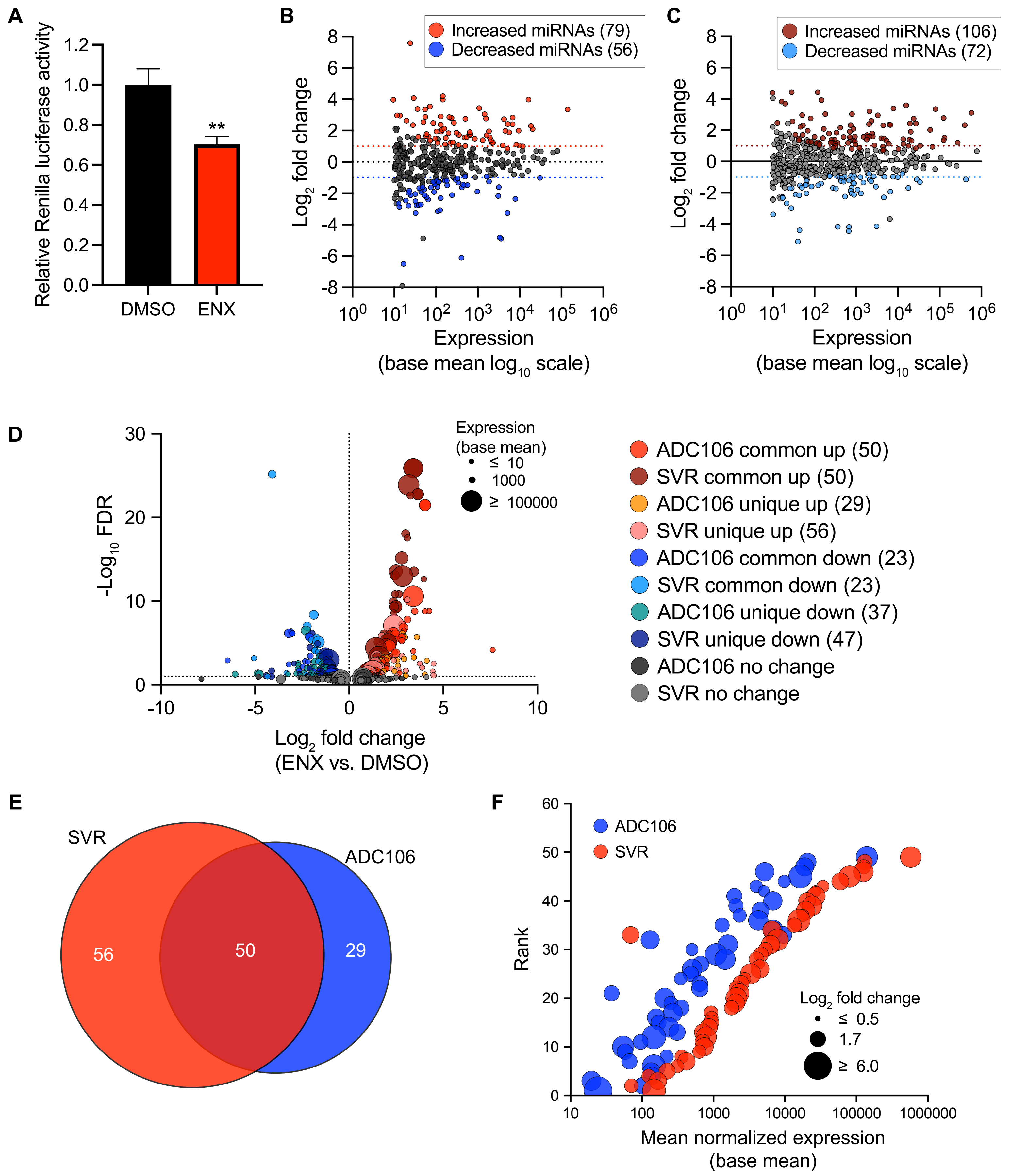
A
Relative Renilla luciferase activity
1.2
1.0
0.8
0.6
0.4
0.2
0.0
**
DMSO
ENX
B
Increased miRNAs (79)
Decreased miRNAs (56)
Log2 fold change
Expression
(base mean log10 scale)
C
Increased miRNAs (106)
Decreased miRNAs (72)
Log2 fold change
Expression
(base mean log10 scale)
D
-Log10 FDR
Expression
(base mean)
≤ 10
1000
≥ 100000
ADC106 common up (50)
SVR common up (50)
ADC106 unique up (29)
SVR unique up (56)
ADC106 common down (23)
SVR common down (23)
ADC106 unique down (37)
SVR unique down (47)
ADC106 no change
SVR no change
Log2 fold change
(ENX vs. DMSO)
E
SVR
ADC106
56
50
29
F
ADC106
SVR
Rank
Log2 fold change
≤ 0.5
1.7
≥ 6.0
Mean normalized expression
(base mean)

Supplementary Figure S2. Murphy *et al*.

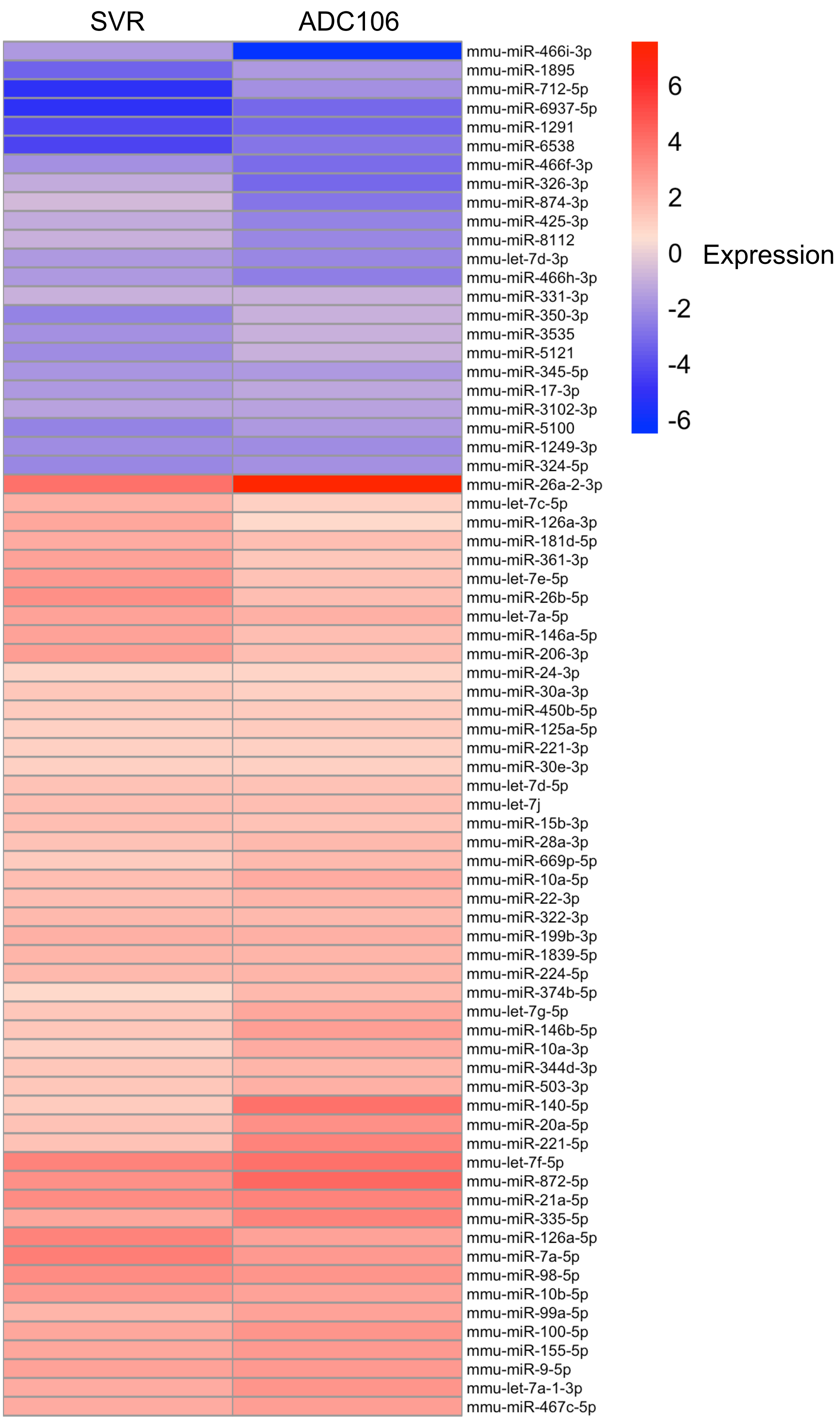

Supplementary Figure S3. Murphy *et al.*

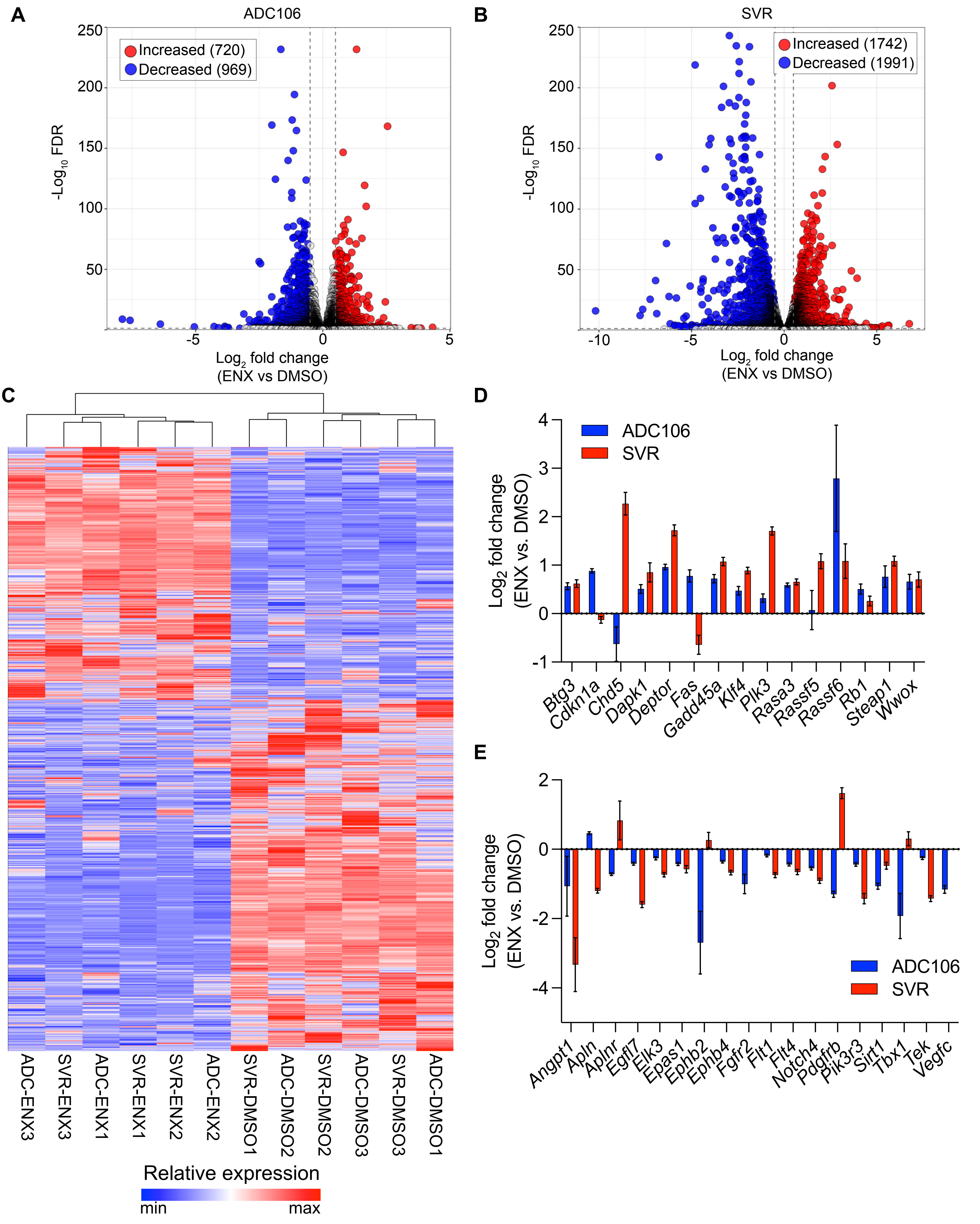

Supplemental Figure S4. Murphy *et al*.

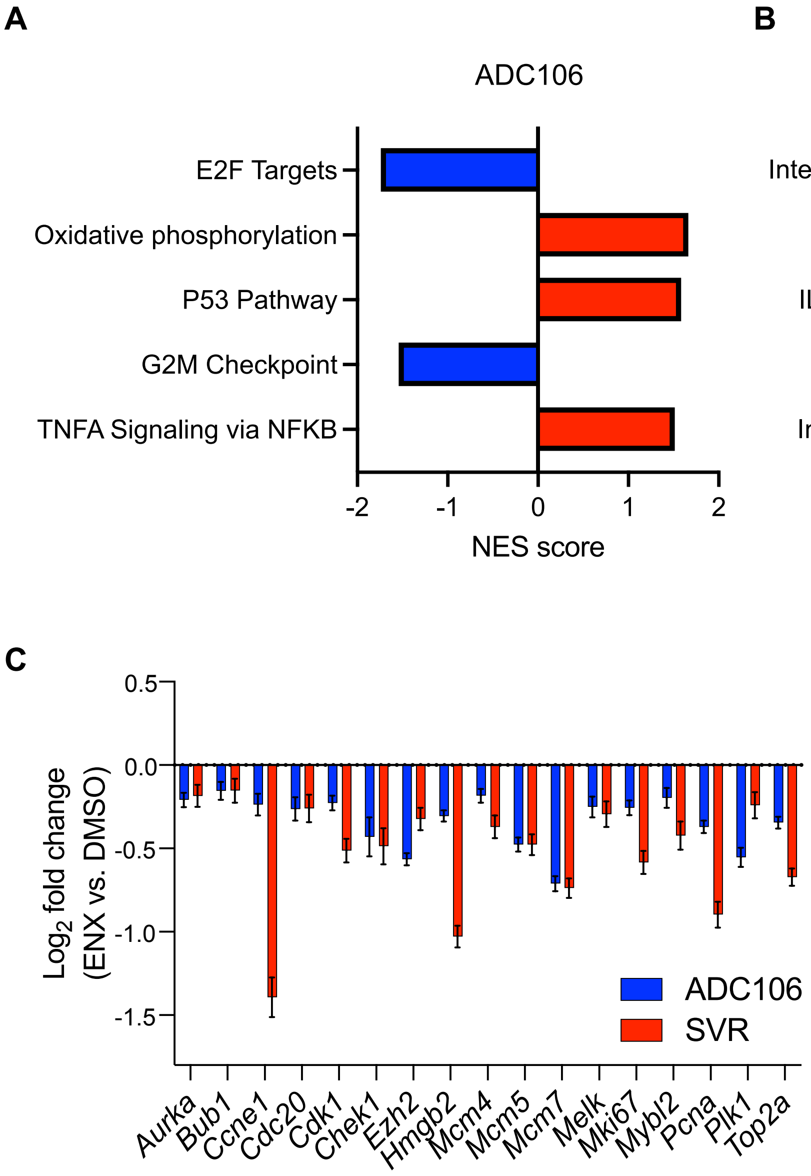
